# Sulcal anatomy of ventral temporal cortex and reading development

**DOI:** 10.64898/2026.04.06.716640

**Authors:** Jewelia K. Yao, Jamie L. Mitchell, Alina Davison, Jason D. Yeatman

## Abstract

Individual differences in cognitive abilities have been linked to variability in cortical folding, a stable neuroanatomical scaffold largely established in utero. In the domain of reading, recent findings in small groups of typical readers suggest that a sulcal interruption (superficial annectant gyrus, gyral gap) in the left posterior occipital temporal sulcus (lhpOTS) predicts better reading skills, posing the lhpOTS as a potential early biomarker of reading difficulties. However, it remains unknown whether this relationship found in typical readers generalizes to the dyslexic population and whether the lhpOTS can serve as a biomarker for dyslexia or predict response to targeted instruction.To fill these gaps, we examine the patterns of the lhpOTS in 209 children, including children with dyslexia, from four independently-collected samples. In typical readers, we find that the relationship between the lhpOTS and reading skills is robust, replicating across binary and continuous quantifications of the sulcal interruption. However, lhpOTS patterns neither distinguish dyslexic children from typical readers nor do they predict response to intervention. Instead, targeted reading intervention drives long-term gains in reading skills that are equivalent irrespective of VOTC anatomy. Together, these findings distinguish neuroanatomical correlates of skilled reading from determinants of reading impairment and learning capacity and emphasize the importance of the educational environment in supporting reading acquisition for children with dyslexia.

**SIGNIFICANCE STATEMENT:** Early predictors of dyslexia are important for understanding the etiology of reading difficulties and informing early intervention. One candidate biomarker for dyslexia is the left posterior occipital temporal sulcus (lhpOTS), a neuroanatomical feature established before birth. In typical readers, the presence of an interruption in the lhpOTS has been linked to better reading skills. Here, we examine this neuroanatomical feature in 209 children with and without dyslexia. While the lhpOTS reliably relates to reading skill in typical readers, it neither differentiates dyslexic from typical readers nor predicts response to intensive reading intervention. These results show that brain anatomy reflects reading proficiency but does not constrain learning and highlights the power of targeted intervention to support reading development.

## INTRODUCTION

A central goal of cognitive neuroscience is to understand how brain anatomy gives rise to cognition. A growing body of work suggests that individual differences in cognitive abilities reflect, in part, variability in cortical folding patterns (Amiez et al., 2018; Chung et al., 2017; Gregory et al., 2016). As cortical folding is established in utero and remains largely stable across the lifespan (Chi et al., 1977; Zilles et al., 2013), sulci and gyri may provide a scaffold for experience-dependent plasticity and the development of cognitive functions (Borst et al., 2016; Bouhali et al., 2024). In humans, sulcal morphology has been associated with individual differences in reasoning (Voorhies et al., 2021; Willbrand et al., 2022, 2023), working memory (Yao et al., 2023), face processing (Parker et al., 2023), inhibitory control (Tissier et al., 2018), and psychopathology (Cachia et al., 2008; Galaburda & Bellugi, 2000; Ramos Benitez et al., 2024). More recently, sulcal interruptions – superficial gyral gaps within a sulcus (annectant gyri; Bodin et al., 2021; Mangin et al., 2019) – have been linked to learned skills like reading (Borst et al., 2016; Bouhali et al., 2024; Cachia et al., 2018), writing (Cachia et al., 2025), math (Roell et al., 2022; Schwizer Ashkenazi et al., 2024), and language (Santacroce et al., 2024), highlighting sulci as features that influence the development of higher order cognitive abilities.

Learning to read is one of the most consequential achievements of childhood, and children vary widely in how efficiently this skill is acquired. For some, reading difficulties lead to a diagnosis of dyslexia, a learning disability characterized by persistent challenges with reading, writing, and spelling. Neuroimaging studies have identified a distributed reading network that includes ventral occipital temporal cortex (VOTC), inferior frontal cortex, and temporal parietal cortex (Dehaene & Cohen, 2011; Pugh et al., 1996; Shaywitz et al., 2002). Relative to typical readers, children with dyslexia exhibit structural (Hoeft et al., 2011; Im et al., 2016; Raschle et al., 2011) and functional (Brem et al., 2020; Kubota et al., 2019; Mitchell et al., 2025; Paulesu et al., 2014; Richlan et al., 2009) differences within this network.

Within VOTC, the occipital temporal sulcus (OTS) has emerged as an important structural landmark for reading. The OTS anchors the location of the visual word form areas (VWFA), functional regions selectively responsive to written words that support fluent word recognition (Dehaene & Cohen, 2011; Grill-Spector & Weiner, 2014; Yeatman et al., 2013). Furthermore, the OTS structure is also linked to reading abilities. Several studies in children with typical reading proficiencies and literate adults (Borst et al., 2016; Bouhali et al., 2024; Cachia et al., 2018, 2025) report that a sulcal interruption in the posterior portion of the left hemisphere OTS (lhpOTS) predicts higher reading abilities, an effect that strengthens across development. Conversely, individuals with a continuous lhpOTS exhibited significantly lower reading scores, posing the lhpOTS as a potential biomarker for reading difficulties.

However, it remains unknown whether the relationship between lhpOTS interruptions and reading generalizes to larger populations with more variable reading abilities, and whether the lhpOTS plays a causal role in shaping reading outcomes. Specifically, no study has explicitly tested whether a continuous lhpOTS is a biomarker for dyslexia. Given evidence that the lhpOTS-reading effect is cumulative (Bouhali et al., 2024), whether the lhpOTS reflects learning capacity and can predict gains from targeted reading instruction is also unknown.

Thus, the present study leverages a large sample of 114 typical readers and 95 dyslexic children from four independently-collected datasets. First, we test if an interrupted lhpOTS predicts better reading abilities in a larger, more heterogeneous population including children with dyslexia. Second, we evaluate whether lhpOTS morphology distinguishes typical readers from dyslexic children, testing its potential as a neuroanatomical biomarker of dyslexia. Finally, in a subset of dyslexic participants (N = 75) who completed an intensive reading intervention, we assess whether lhpOTS morphology predicts responsiveness to instruction, isolating its potential influence on learning by controlling the educational environment.

## MATERIALS AND METHODS

### Participants

#### Cross-sectional Participants

To obtain a large sample size with greater variability in reading abilities, we leverage previously published data from four independent studies (**Table 1**): the Stanford Intervention (Stanford-I) study (N = 84; Mitchell et al., 2025), the Seattle Intervention (Seattle-I) study (N = 46; Huber et al., 2018), the Seattle Developmental (Seattle-D) study (N = 47; (Caffarra et al., 2024; Yeatman et al., 2024), and the Stanford Developmental (Stanford-D) study (N = 32; Nordt et al., 2019). Across studies, 209 participants (ages 5 - 13; 102 females) were included in the present cross-sectional analyses. The Stanford-D sample was recruited with the intention of including only typical readers, but the Stanford-I, Seattle-I, Seattle-D samples were recruited with the intention to also include participants with dyslexia. The Seattle-D study includes children who have not started formal schooling. All participants reported normal or normal to corrected vision, no history of psychiatric disorder or neurological damage, and were native speakers of English. The study protocols for the Stanford-I and Stanford-D studies were approved by the Stanford Internal Review Board on Human Subjects Research. The study protocols for the Seattle-I and Seattle-D studies followed the guidelines of the University of Washington Human Subjects Division and were approved by the University of Washington Institutional Review Board. All participants and their parents gave their informed assent and consent.

**Table 1.** Participant characteristics across four independent studies. N, age, TOWRE Composite Index scores, and diagnostic breakdown for each study included in cross-sectional analyses. Dyslexia classification was defined as TOWRE Index < 85; typical readers had TOWRE Index ≥ 85. Stanford-D was recruited as a typical reader sample; Stanford-I, Seattle-I, and Seattle-D were recruited to include participants with and without dyslexia. Seattle-D includes children who had not yet begun formal schooling. M = mean; SD = standard deviation.

| Study | N | Age: $M \pm SD$ | TOWRE Index: $M \pm SD$ | Dyslexic | Typical |
| --- | --- | --- | --- | --- | --- |
| Stanford-I | 84 | $10.0 \pm 1.4$ (7.4–13.8) | $80.8 \pm 14.1$ (56–125) | 54 | 30 |
| Seattle-I | 46 | $9.5 \pm 1.6$ (7.2–12.7) | $79.7 \pm 18.6$ (53–127) | 30 | 16 |
| Seattle-D | 47 | $6.2 \pm 0.3$ (5.7–7.0) | $95.3 \pm 13.6$ (72–134) | 10 | 37 |
| Stanford-D | 32 | $8.3 \pm 2.3$ (5.0–12.0) | $131.6 \pm 19.5$ (78–147) | 1 | 31 |

#### Longitudinal Participants

Longitudinal data were collected from participants in the Stanford-I and Seattle-I studies. 42 participants from the Stanford-I study and 29 participants from the Seattle-I study participated in an intensive summer reading intervention program (Bell, 2013). Members of the intervention groups were recruited based on clinical diagnosis of dyslexia and/or parent report of reading difficulties. Forty-two participants from the Stanford-I study and 22 participants from the Seattle-I study acted as controls for each study. The control groups included both typical readers and dyslexic readers who did not receive the intervention program but were measured at the same timepoints as the intervention participants. In the Stanford-I study, behavioral and MRI sessions occurred at five timepoints: baseline, pre-intervention, post-intervention, a 6 month follow-up, and a 12-month follow up. 40 participants completed all five timepoints, 68 participants completed at least four timepoints, 71 completed at least three timepoints, and 83 completed at least two timepoints. In the Seattle-I study, behavioral and MRI sessions occurred at four timepoints: pre-intervention, after 2.5 weeks of the intervention, after 5 weeks of the intervention, and at the end (8 weeks) of the intervention. 23 participants completed all four timepoints, 28 completed at least three timepoints, and 29 completed at least two timepoints. For the comparison across Stanford-I and Seattle-I studies, our analysis focuses on the pre-intervention and post-intervention timepoints from each dataset. Participants with only one timepoint from the Stanford-I and Seattle-I studies and participants from the Seattle-D and Stanford-D studies, which only had one timepoint, were not included in the longitudinal analysis.

#### Reading Intervention

Intervention participants from the Stanford-I and Seattle-I studies were enrolled in 8 weeks (4 hours a day, 5 days a week, 8 weeks) of the Seeing Stars: Symbol Imagery for Fluency, Orthography, Sight Words, and Spelling program at Lindamood-Bell Learning Centers in the Bay Area (Stanford-I) or Seattle area (Seattle-I) (Bell, 2013). The intervention program included directed, personal one-on-one training in phonological and orthographic processing skills. The curriculum used an incremental approach, building from letters and symbols to words and text, emphasizing phonological decoding as the foundation for spelling and comprehension. Due to the COVID-19 pandemic, intervention sessions for children in the Stanford-I study were conducted remotely.

#### Dyslexia Grouping

As some children with reading difficulties do not have an official dyslexia diagnosis and because diagnosis criterion varies among practitioners (Siegel, 2006), we utilized the Test of Word Reading Efficiency (TOWRE; Torgesen et al., 2012) Index, described below, to group participants into typical readers (N = 114; Stanford-I: 29, Seattle-I: 14, Seattle-D: 37, Stanford-D: 32), those with TOWRE Index scores ≥ 85, and dyslexic readers (N = 95; Stanford-I: 55, Seattle-I: 32, Seattle-D: 10, Stanford-D: 0), those with TOWRE Index scores < 85, at the first timepoint of each study. The TOWRE Index is an age-normed, standardized measure of reading skill. The cut-off of 85 indicates one standard deviation below the population mean and is frequently used as a criterion for defining dyslexia (Rimrodt et al., 2009; Shaywitz et al., 2002).

### Experimental Design and Statistical Analyses

#### Reading Measures

Reading measures were collected at each timepoint for the Stanford-I and Seattle-I studies and at a single timepoint for the Seattle-D and Stanford-D studies. On the same day as the MRI session, participants completed a series of behavioral tests. Across all studies, reading scores were measured using the Test of Word Reading Efficiency (TOWRE) subtests for sight word efficiency (SWE) and phonemic decoding efficiency (PDE). These subtests measured the number of sight words (SWE) or pseudowords (PDE) correctly read aloud in 45 seconds. The TOWRE Index is a composite score of the SWE and PDE adjusted for age. We used TOWRE Index as the primary measure of reading skill because it is comparable to or the same as the verbal and speed-based tests used in previous studies examining OTS morphology and reading abilities (Borst et al., 2016; Bouhali et al., 2024; Cachia et al., 2018), allowed us to compare across a range of ages, and was administered in all included studies. Due to the COVID-19 pandemic, assessments for the Stanford-I participants were administered through Zoom.

#### MRI Data Acquisition and Processing

T1-weighted scans were collected at multiple sites with different protocols, listed below. In all studies, T1-weighted images were visually inspected for artifacts. Using FreeSurfer’s automated segmentation tools (http://surfer.nmr.mgh.harvard.edu; Dale et al., 1999), each anatomical volume was segmented to separate white and gray matter, and the resulting boundary was used to reconstruct the cortical surface for each subject. Each cortical reconstruction was inspected for segmentation errors and manually corrected when necessary.

### Stanford-I

MRI data for the Stanford-I study were collected at the Stanford University Center for Cognitive and Neurobiological Imaging (CNI) using a 3T Signa GE Scanner with a 32-channel coil. Whole-brain, high-resolution T1-weighted anatomical scans were acquired with 0.9mm3 isotropic voxels. The T1w image was reoriented into AC–PC alignment with ANTSpy version 0.4.2 (Tustison et al., 2021), and then skull-stripped using *Synthstrip* (Hoopes et al., 2022). Segmentation and cortical reconstruction were then performed using *Synthseg* robust algorithm (Billot et al., 2023) for segmentation and surface reconstruction, as implemented in FreeSurfer 7.3.2.

### Seattle-I and Seattle-D

MRI data for the Seattle-I study were collected at the the University of Washington Diagnostic Imaging Sciences Center (DISC) using a 3T Phillips Achieva scanner with a 32-channel head coil. Whole-brain, high-resolution T1-weighted MPRAGE anatomical scans were acquired with 0.8mm3 isotropic voxels. Segmentation and cortical reconstruction were performed using FreeSurfer 7.3.2.

### Stanford-D

MRI data for the Stanford-D study were collected at the Stanford University CNI using a 3T Signa GE Scanner with a 32-channel coil. Whole-brain, high-resolution anatomical scans were acquired using T1-weighted quantitative MRI (qMRI, Mezer et al., 2013), using a spoiled gradient echo sequence with multiple flip angles (α = 4°,10°,20°, and 30°;TR= 14 ms; TE = 2.4 ms). Voxel size = 0.8 mm × 0.8 mm × 1 mm, resampled to 1 mm3 isotropic. An artificial T1-weighted anatomy was generated from qMRI data using mrQ (https://github.com/mezera/mrQ). Segmentation and cortical reconstruction were performed using FreeSurfer 5.3.0.

### Morphological Analyses

#### Sulcal definitions

The OTS and other VTC sulci were identified manually for each participant at the first timepoint in each study. Sulci were separately defined by JKY and AL and corrected if disagreement occurred. Before corrections, inter-rater reliability was ∼90 % across all sulci. Labeling was conducted blind to dyslexia diagnosis and reading scores. Sulci were defined on the inflated surface generated from FreeSurfer, with cross-referencing to folding patterns observed in the pial and white surfaces, also generated from Freesurfer.

OTS definitions were based on those used in previous studies linking reading and sulcal patterns ((Borst et al., 2016; Bouhali et al., 2024; Cachia et al., 2018). To define the OTS, we first identified surrounding VOTC sulci (MFS: mid fusiform sulcus, CoS: collateral sulcus, atCoS: anterior transverse CoS, ptCoS: posterior transverse CoS) and the pre-occipital notch, the boundary between occipital and temporal cortices, based on previously established definitions (Duvernoy & Vannson, 1999; Petrides, 2019; Weiner et al., 2014)). The posterior VTC boundary was defined by a virtual line between the ptCoS and pre-occipital notch, which also acted as the posterior boundary of the OTS; any sulcal component below this line was not considered in our definition of the OTS. The anterior endpoint of the OTS varied by subject, but often met with or ended near the anterior tip of the CoS or the atCoS. Medially, we defined the MFS to distinguish OTS components. Laterally, we also identified the ITS and STS to distinguish OTS components. VOTC sulcal definitions for all subjects can be found in **Fig. S1**.

#### OTS interruption definitions

OTS interruptions were defined by manually defining gyral crowns between OTS sulcal components. An interruption was considered posterior (pOTS) if it was located below the anterior tip of the MFS and anterior (aOTS) if it was located above the anterior tip of the MFS. The anterior tip of the MFS approximates MNI Y=-40 and was used as the anterior-posterior boundary in Bouhali et al. (2024). As segmentation errors may lead to artificial interruptions in the OTS, we only considered an interruption present when its size was greater than 10mm in the case of our binary interruption measure. OTS, lhpOTS interruption, and lhaOTS interruption definitions for all subjects can be found in **Fig. S2**.

We quantified the OTS interruptions in three ways: (i) interruption presence, a binary measure where having an interruption greater than 10mm in length was considered present and one less than 10mm was considered not present, (ii) interruption length, the geodesic distance in mm from the most posterior to most anterior point in the interruption, and (iii) interruption ratio, the number of voxels in an interruption over the total number of voxels in the sulcal components and interruption components (aOTS + pOTS). While we quantified these measures for the aOTS and pOTS, our main analysis only includes the pOTS interruption because only the left pOTS has been linked to reading abilities in prior studies (Borst et al., 2016; Bouhali et al., 2024; Cachia et al., 2018).

#### Sulcal measurements

Additionally, we calculated the geodesic distance (length), cortical thickness, average sulcal depth, and maximum sulcal depth for the OTS and all other sulci (MFS, CoS, atCoS, ptCoS) for comparison using custom code (https://github.com/yeatmanlab/OTS-Anatomy-Dyslexia) extracting and calculating measures from FreeSurfer sulc values.

### Statistical Tests

#### Cross-Sectional analysis

To test the hypothesis that the lhpOTS interruption predicted reading abilities, we compared three nested linear models predicting TOWRE Index using the lm package in R:

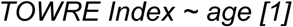

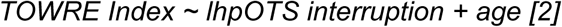

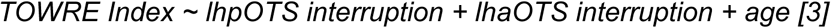

Age was included in all models because it correlates with *TOWRE* Index scores. Improvement in fit was evaluated using likelihood ratio tests (ANOVA). This set of models was used to evaluate the relationship between lhpOTS interruption and reading abilities separately for the binary interruption, interruption length, and interruption ratio measures.

To determine if the link between lhpOTS interruption and reading differed by diagnosis, we fit the linear model *TOWRE Index ∼ diagnosis * lhpOTS interruption + age*. A significant interaction was followed by simple effects tests within typical and dyslexic subgroups (ANCOVA, controlling for age).

#### Longitudinal analysis

Linear models were fitted using the lm package in R. To determine if sulcal patterns predicted reading abilities at the end of the intervention, we first fit a model *TOWRE Index Post-Intervention ∼ lhpOTS interruption* + *age (at time of T1 used for sulcal definition)* within the intervention group. We ran additional analysis with this model for different timepoints throughout the intervention period and at the follow-up. To determine if sulcal patterns predicted growth, we fit a linear model predicting the difference between pre- and post-intervention TOWRE Indices from the same variables (*Post - Pre TOWRE Index ∼ lhpOTS interruption + age*). To compare the effects of anatomy versus the intervention on scores, we fit a linear model that additionally included whether a participant was enrolled in the intervention or not (*TOWRE Index Post-Intervention ∼ lhpOTS interruption + intervention + age*).

### Code Accessibility

Custom code for extracting morphological measures were based on code from (Voorhies et al., 2021; Yao et al., 2023). Code and processed data for reproducing figures and statistics can be found at https://github.com/yeatmanlab/OTS-Anatomy-Dyslexia. Raw data available upon request.

## Results

### An interrupted left posterior OTS is associated with better reading skills across four independent sample

The anatomy of the OTS varies dramatically. In some brains, the OTS stretches continuously from the posterior transverse collateral sulcus to the anterior end of the collateral sulcus, whereas in other brains the OTS is interrupted by small gyri that break the OTS into 2 or 3 segments. To test if the morphology of the lhpOTS relates to reading scores in a larger cohort with more variability in reading ability (including those with dyslexia), we manually delineated the OTS and its interruptions (**Fig. 1A**; lhpOTS, lhaOTS) in both hemispheres across four independent samples of children (N = 209; see Methods). For subsamples with longitudinal data, only the first timepoint was included since sulcal patterns are stable after birth (Chi et al., 1977; Meng et al., 2014; White et al., 2010). For comparison, we defined the MFS, CoS, atCoS, and ptCoS as well. All sulcal and gyral definitions were conducted blinded to dyslexia diagnosis and reading scores.

**Figure 1.**
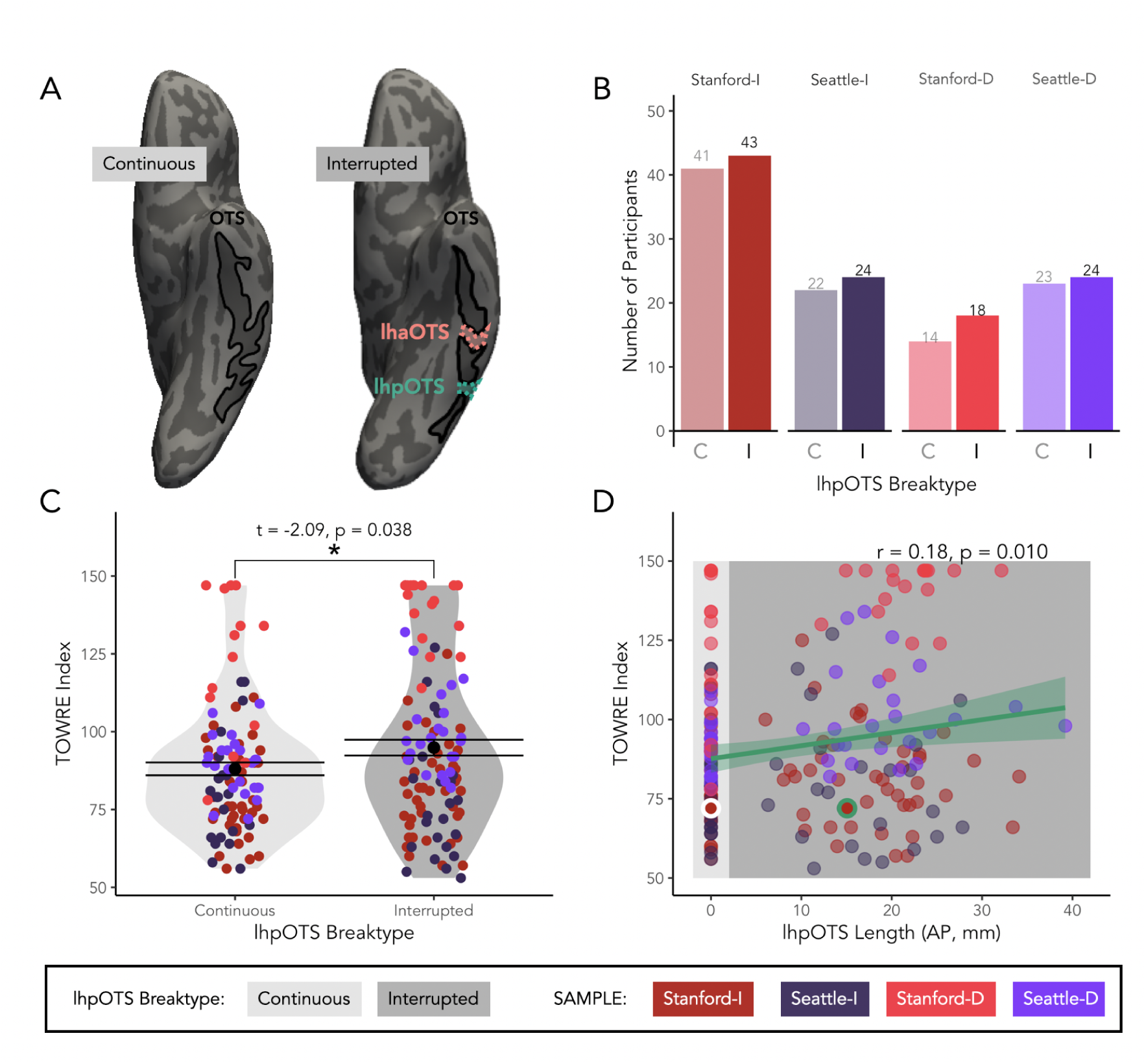
Replication of the relationship between the lhpOTS interruption and reading abilities. A. *Left:* Inflated view of left hemisphere ventral temporal cortex with a continuous OTS (black outline) in an example participant (9 years of age). *Right:* Inflated view of left hemisphere ventral temporal cortex with an interrupted OTS (black outline) in a different participant (9 years of age). Coral dotted outline: lhaOTS interruption. Turquoise dotted outline: lhpOTS interruption. B. Histogram of occurrence of continuous (C, light shaded bars) and interrupted (I, dark shaded bars) left hemisphere pOTS interruptions in the Stanford Intervention Study (Stanford-I; dark red), the Seattle Intervention Study (Seattle-I; dark purple), the Stanford Developmental Study (Stanford-D; red), and the Seattle Developmental Study (Seattle-D; purple). C. Violin plots of TOWRE Index scores for children with continuous (light gray) versus interrupted (dark gray) lhpOTS. Each dot, colored by sample, is an individual participant. Asterisk indicates a significant difference between the two groups. D. Correlation between lhpOTS interruption ratio and TOWRE Index for the same children. Each dot, colored by sample, is an individual child. Light gray shaded box: children with continuous OTS. Dark gray shaded box: children with an interrupted OTS.

Across all samples, we observed a high prevalence of OTS interruptions consistent with previous studies (**Fig. 1B**). Overall, the OTS showed at least one interruption in 88.5% of left hemispheres and 82.3% of right hemispheres (**Fig. S1**). The lhpOTS was interrupted in 52.2% of children, and the rhpOTS was interrupted in 45% of children. Between samples, we observed no significant differences in the occurrence of lhpOTS interruptions (X^2^(3, N = 209) = 0.27, p = 0.97). The anterior left OTS (lhaOTS) was interrupted in 77% of children, and the rhaOTS was interrupted in 67.9% of children. We observed no significant differences in the occurrence of lhaOTS interruptions between samples (X^2^(3, N = 209) = 0.86, p = 0.83). The occurrence of the lhpOTS was independent from the occurrence of lhaOTS (X^2^(1, N = 209) = 0.03, p = 0.86).

To assess whether lhpOTS interruptions were associated with better reading performance, we used an ANOVA to compare three nested linear regression models predicting TOWRE Index by age [1], by age and presence of an lhpOTS interruption (binary predictor) [2], and by age, lhpOTS interruption presence, and lhaOTS interruption presence [3]. The model including lhpOTS interruption presence [2] was significantly better at predicting TOWRE Index scores, above and beyond a model with age alone [1] (F(1, 206) = 4.1985, p = 0.04173), though the effect was small. Importantly, children with an interrupted lhpOTS (M = 94.88 ± 26.49) had significantly higher reading scores than those with a continuous lhpOTS (**Fig. 1C**; M = 88.05 ± 20.68297; t (201.94) = -2.0867, p = 0.03817), replicating previous findings. Interestingly, including the lhaOTS interruption in the model [3] did not significantly improve model fit (F(1, 205) = 1.8193, p = 0.1789). In fact, the presence of the lhaOTS interruption did not predict scores in the combined model [3] or in a model including only age and the lhaOTS (p’s > .05). In sum, we successfully replicate previous methods and findings in a large, multi-site sample and find that the binary presence of the lhpOTS is associated with reading scores in the overall population.

### Continuous measures of both left OTS interruptions strongly predict reading abilities

While the binary presence of an interruption captures whether the OTS is segmented versus continuous, it does not reflect how much of the sulcus is interrupted (i.e., the size of the annectant gyrus) nor does it account for individual differences in sulcal size. To quantify the degree of sulcal segmentation, we calculated two continuous interruption measures: 1) the anterior-posterior length of the lhpOTS interruption, or the geodesic distance between the most anterior and posterior vertices of the interruption (**Fig. 1D**), and 2) the interruption ratio, the number of vertices in the interruption over the total number of vertices in all OTS interruptions and sulcal components, which accounts for individual variation in size of the OTS (**Fig. 2**).

**Figure 2.**
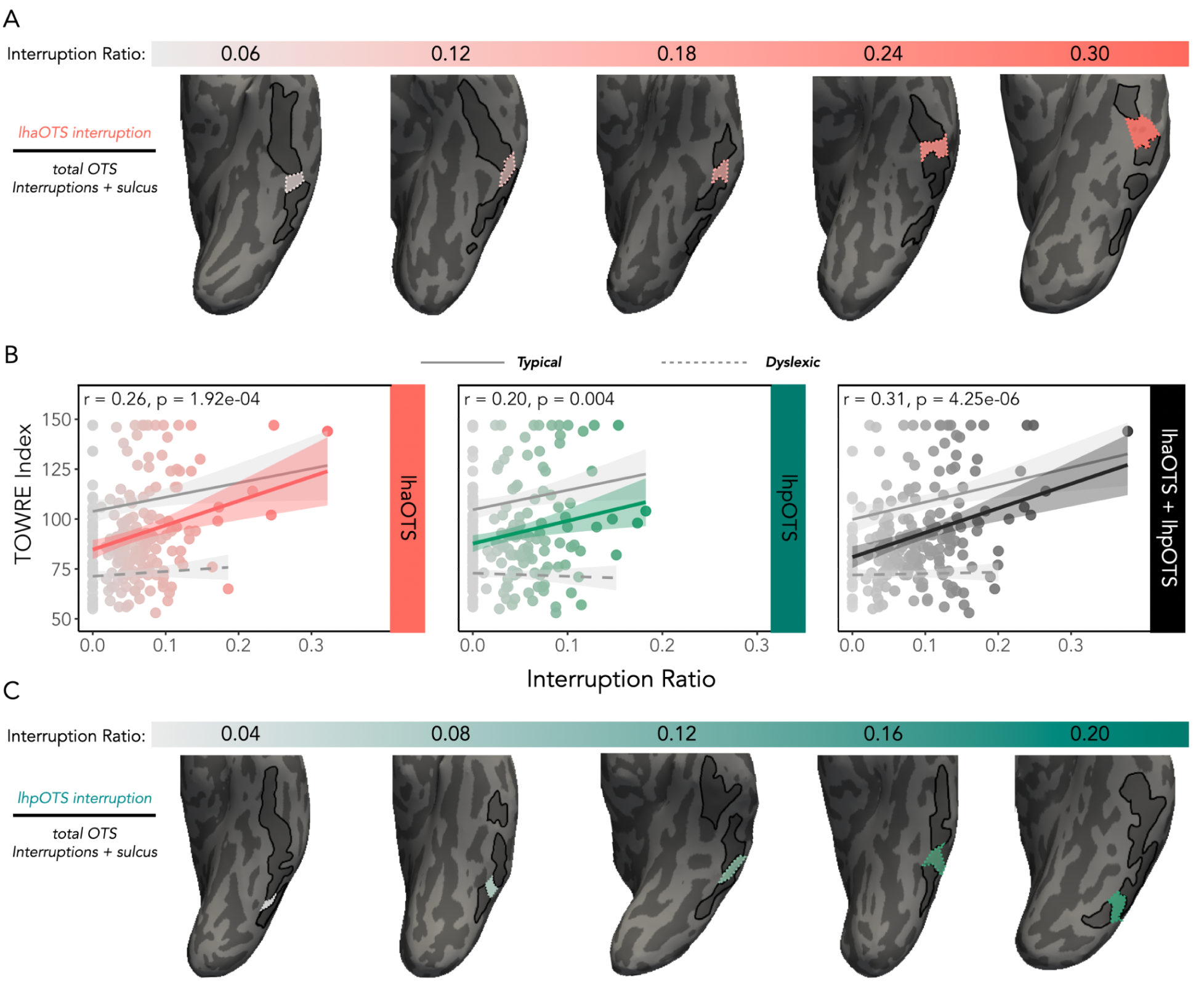
Gap ratio of the lhpOTS and lhaOTS and its relationship to reading abilities. A. Example of the lhaOTS interruption (coral) with lower interruption ratios on the left (light gray) transitioning to higher interruption ratios on the right (dark coral) in five example subjects. B. *Left:* relationship between lhaOTS (coral) interruption ratio and TOWRE Index for all subjects. Light gray dot: individual with no interruption. Coral dot: individual with interruption; higher saturation = higher interruption ratio. Coral line: linear relationship between interruption ratio and TOWRE Index for all subjects. Gray solid line: linear relationship between interruption ratio and TOWRE Index for typical readers (all significant: r’s > 0.2, p’s < .031); Gray dotted line: linear relationship between interruption ratio and TOWRE Index for dyslexic children, all n.s. *Middle:* Same as left but for lhpOTS (turquoise). *Right:* Same as left but for the combined ratio of lhpOTS and lhaOTS interruptions. C. Example of the lhpOTS interruption (turquoise) with lower interruption ratios on the left (light gray) transitioning to higher interruption ratios on the right (dark turquoise) in five example subjects.

We examined the relationship between TOWRE Index scores and these lhpOTS continuous measures, controlling for age, using linear regression models (*TOWRE Index ∼ age + lhpOTS length or interruption ratio*). Both lhpOTS length and interruption ratio were significant predictors of reading performance, such that children with longer or larger interruptions in the sulcus exhibited higher reading scores. Compared to the binary lhpOTS measure [2] (*ß* = 6.6685, p = 0.042), both continuous measures provided stronger predictions of reading ability, with lhpOTS interruption ratio emerging as the best single predictor (*ß* = 105.5891, p = 0.0076; **Fig. 2A,B**), followed by lhpOTS interruption length (*ß* = 0.3865, p = 0.0146; **Fig. 1D**).

Additionally, as the lhaOTS interruption is present in most children, we reasoned that a continuous measure might be more sensitive to individual differences in lhaOTS morphology and its relationship to behavior. We therefore also examined whether continuous metrics of lhaOTS interruptions were also related to reading ability. Indeed, in models where the lhaOTS length or interruption ratio was included, both lhaOTS interruption length (*ß* = 0.3832, p = 0.01201) and lhaOTS interruption ratio (*ß* = 125.332, p = 8.27e-05) were significant predictors of reading scores. Unlike the binary interruption measures, continuous metrics revealed independent, additive contributions of anterior and posterior OTS interruptions to reading abilities with the combined interruption ratio model (**Fig. 2B,C**), compared to other models and interruption measures, explaining the greatest variance in TOWRE Index scores (Adj-R^2^ = 0.122, p = 0.0000015).

### Anatomy-behavior relationship is specific to the left OTS

We next conducted an exploratory analysis to determine if the reported effects were a) specific to the OTS versus b) reflective of broader neuroanatomical differences across VOTC. Importantly, we found that the relationship between reading ability and sulcal morphology was specific to the OTS interruptions themselves. To explore the effect of other VOTC sulci on reading performance, we fit an exploratory analysis correlating morphological measures (length, mean, depth, cortical thickness) of sulcal components of the OTS and neighboring ventral occipitotemporal sulci (CoS, atCoS, ptCoS, mfs) with TOWRE Index. No correlations were significant before and after correction for multiple comparisons (BH; p’s > .05). Thus, the relationship between reading performance and cortical morphology appears localized to the OTS rather than reflecting differences in VOTC folding patterns more broadly.

### A continuous lhpOTS is not a biomarker of dyslexia

Previous studies proposed that the lhpOTS interruption could serve as a potential biomarker of dyslexia (Borst et al., 2016; Bouhali et al., 2024; Cachia et al., 2018). However, earlier studies included very few (if any) participants with dyslexia. Next, we specifically test whether a) the association between OTS anatomy and reading is present in both children with dyslexia and typical readers and b) the lhpOTS can be used to predict learning differences in responses to a reading intervention.

Although lhpOTS patterns predict reading scores across the full sample, we tested whether this relationship extends to dyslexic children and whether a continuous lhpOTS is a biomarker for developmental dyslexia. In our samples, 95 children were classified as dyslexic (TOWRE Index < 85) and 114 as typical readers (TOWRE Index ≥ 85). Notably, both groups exhibited variability in lhpOTS patterns: lhpOTS interruptions were present in 62% of typical readers and 47% of dyslexic children (**Fig. 3A**). Qualitatively, there was large overlap in the average lhpOTS interruption locations (**Fig. 3B**), and the difference in interruption occurrence between groups was not significant (X^2^(1, N = 209) = 1.0034, p = 0.3165), demonstrating that dyslexia is not uniformly characterized by a continuous lhpOTS.

**Figure 3.**
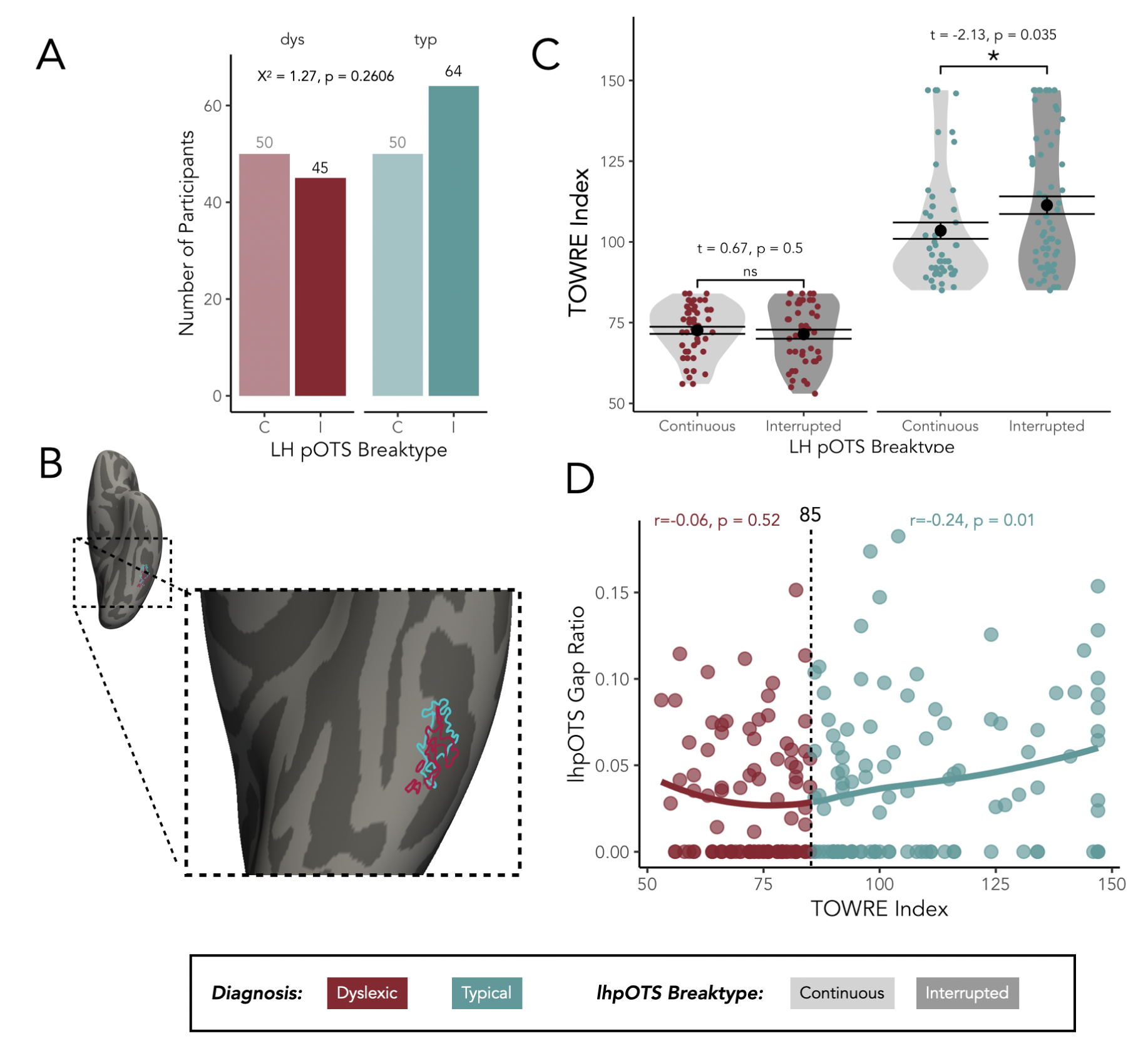
lhpOTS in children with dyslexia versus typical readers. A. Histogram of occurrence of continuous (C, light shaded bars) and interrupted (I, dark shaded bars) left hemisphere pOTS interruptions in children with dyslexia (maroon) and typical readers (teal). B. Probability maps (30% threshold) of the lhpOTS in children with dyslexia and typical readers projected onto the inflated cortical surface. C. Violin plots of TOWRE Index scores for dyslexic children and typical readers with continuous (light gray) versus interrupted (dark gray) lhpOTS. Each dot, colored by diagnosis, is an individual participant. Asterisk indicates a significant difference between the two groups. (D) Loess curve of lhpOTS interruption ratio by TOWRE Index. Each dot represents a participant, colored by either dyslexia (red) or typical reader (blue). Dashed line indicates the dyslexia cutoff.

Next, we examined if the relationship between lhpOTS patterns and reading scores differed between typical readers and dyslexic children using a linear model with lhpOTS binary interruption presence, diagnosis, age, and their interactions as predictors of TOWRE Index. As expected, diagnosis predicted reading scores (F(1,204) = 249.374, p < 0.0001), with typical readers outperforming (M = 108.53 ± 20.325) dyslexic children (M = 72.44898 ± 8.713211). Interestingly, the lhpOTS did not predict reading scores in this model (F(1,204) = 2.911, p = 0.0895), but we found that the relationship between lhpOTS and reading scores differed by diagnosis (diagnosis x lhpOTS: (F(1,204) = 4.526, p = 0.0346).

To explore this interaction, we examined the relationship between lhpOTS interruption and TOWRE scores by group (**Fig. 3C**). Like in previous studies, the lhpOTS binary presence significantly predicted reading abilities for typical readers, whereby children with an interrupted lhpOTS have higher TOWRE scores (F(1,108) = 5.251, p = 0.0239). However, among dyslexic children, lhpOTS interruption presence was not associated with reading performance (F(1,95) = 0.024, p = 0.8779). Thus, while lhpOTS morphology relates to reading abilities in typical development, it does not differentiate dyslexic from typical readers and is not a reliable biomarker for predicting dyslexia diagnosis.

### The lhpOTS does not predict intervention response in children with dyslexia

Prior work in typical readers finds that the interruption effect is cumulative (Bouhali et al., 2024): the strength of the relationship between an lhpOTS interruption and reading scores increases over time. Thus, it is possible that dyslexic children with an interruption in the lhpOTS have better reading skills, or growth, long-term. Conversely, if a continuous OTS is a risk factor for challenges in learning to read, then we would expect to see differences in intervention-driven learning rates for those with an interrupted versus continuous OTS. To test this hypothesis, we leveraged two subsamples of our cohort with longitudinal data collected over the course of an intensive reading intervention study. In one sample (Stanford-I, Fig. 4 - left), 38 dyslexic children took part in an 8 week, intensive (4 hours a day, 5 days a week), one-on-one, intervention, and 18 dyslexic children served as non-intervention controls. In another sample (Seattle-I, Fig. 4 - right), 19 dyslexic participants took part in the same intensive intervention program with anatomical and behavioral data collected at 4 timepoints spaced equally throughout the intervention period. Anatomical and behavioral data were collected at the start and end of the intervention and at two follow-up timepoints (see Methods).

**Figure 4.**
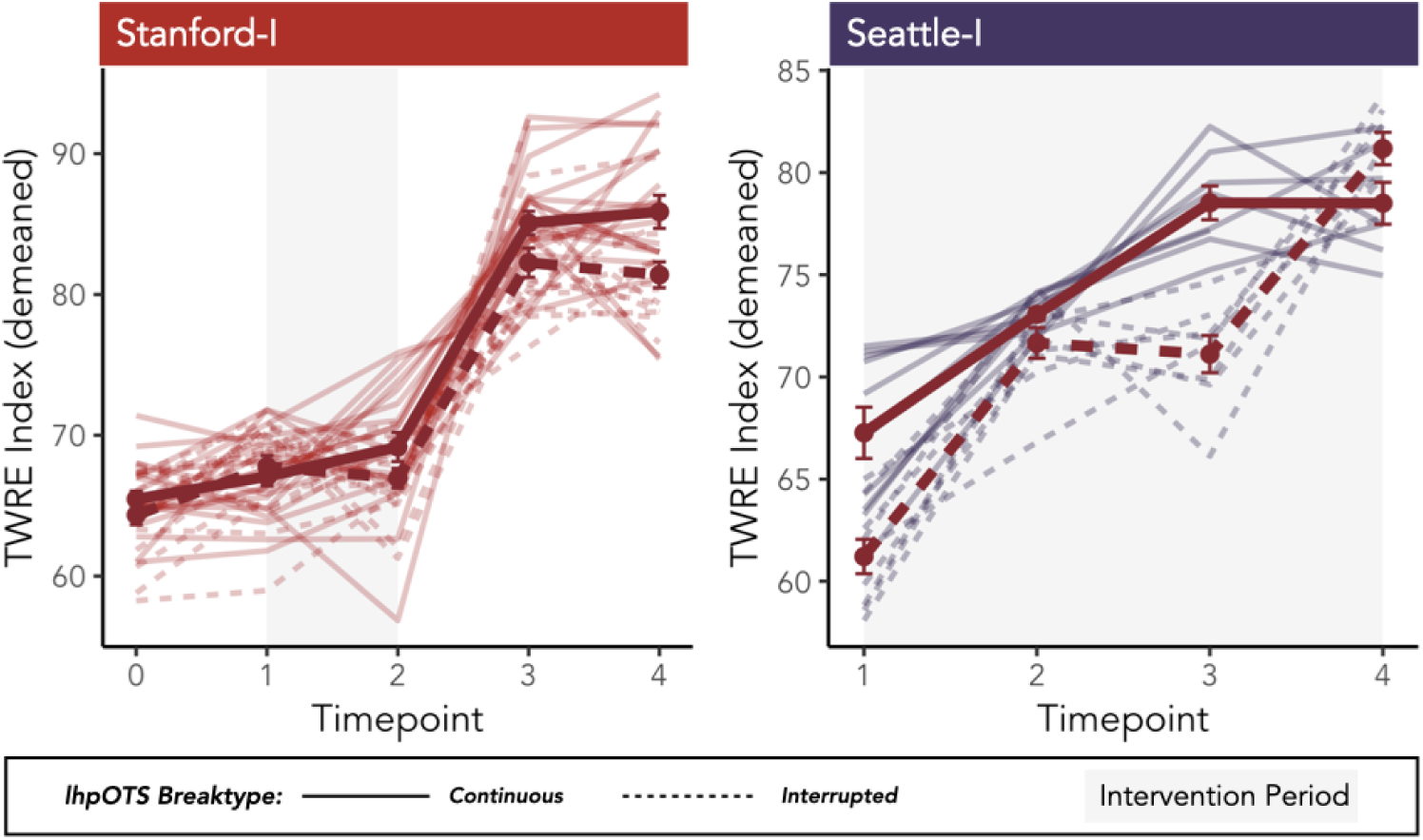
Reading scores across the intervention in dyslexic children with continuous and interrupted lhpOTS. *Left:* Demeaned lineplots of TOWRE Index scores in children with continuous (solid lines) and interrupted (dashed lines) lhpOTS across measurements taken at baseline (0), pre-intervention (1), post-intervention (2), 6-months follow-up (3), and 12-months follow-up (4) from the Stanford Intervention Study (Stanford-I). Light red lines: individual children. Red lines: average scores across time. Light gray shaded box indicates the 8 week intervention period. *Right:* Same as left but for children from the Seattle Intervention Study (Seattle-I). Light purple lines: individual children. Red lines: average scores across time. Measurements were taken at ∼two week intervals during the intervention (light gray shaded box).

For children in the intervention group, we first examined whether having an lhpOTS interruption predicted higher reading scores at subsequent time points or predicted growth rates across the intervention period. A linear model predicting TOWRE Index at the end of the intervention from age and lhpOTS interruption presence finds no significant main or interaction effects of lhpOTS pattern on reading scores (F’s <1.04, p’s > 0.31). Furthermore, a linear mixed-effects model predicting growth rate (slope of line of score change between pre and post-intervention) from age and interruption presence (random intercepts by participant) showed no effect of lhpOTS interruption on the difference in scores from pre-intervention to post-intervention timepoints (F’s < 0.049, p > 0.8257). Thus, when learning opportunities were equated (i.e., everyone received the same curriculum and dosage of reading instruction), anatomical variation in lhpOTS did not predict who benefited most from intervention. In fact, compared to dyslexic controls, children who took part in the intervention exhibited significantly larger improvements in TOWRE scores over the same timeframe (intervention effect: F(1,67) = 14.244, p = 0.0003). Taken together, these results indicate that the anatomical predictors identified in typical readers are not diagnostic in dyslexic populations and highlight how educational experiences play a critical role in shaping reading outcomes in dyslexia.

## Discussion

We examined whether the anatomy of the OTS relates to reading ability and whether the presence of a sulcal interruption constitutes a biomarker for dyslexia. Across four independent samples, we replicate prior reports that the presence of a gyrus interrupting the lhpOTS is associated with better reading performance, particularly among typical readers. Importantly, continuous metrics capturing the size of the interruptions account for more variance in reading skill than a binary classification. Moreover, we show that features of both the posterior and anterior OTS make independent contributions to reading ability. Despite these associations, OTS anatomy neither differentiates dyslexic from typical readers nor predicts reading gains following an intensive dyslexia intervention. When children with dyslexia are provided a high-quality educational experience, learning rates are equivalent regardless of OTS morphology. Together, these findings dissociate anatomical correlates of reading proficiency from biomarkers of dyslexia and from constraints on learning, clarify the role of anatomical variation in VOTC in reading and dyslexia, and underscore the profound effect of evidence-based intervention for all students irrespective of their unique VOTC anatomy.

### A reproducible link between the OTS and reading emerges from stable yet variable folding patterns

Replication in a large sample of four independent studies and across different metrics of sulcal interruptions extends prior findings in smaller, homogeneous cohorts and demonstrates that the relationship between OTS morphology and reading is robust and generalizable. The prevalence and spatial consistency of sulcal interruptions provide a stable structural basis for linking the brain to behavior. Classic work (Cunningham, 1890; Regis, 1994) characterizes these interruptions – annectant gyri – as frequent and systematically organized features of cortical folding that emerge early in development. Both lhpOTS and lhaOTS interruptions have been documented in neuroanatomical atlases (Ono et al., 1990; Petrides, 2019) and in empirical studies (Borst et al., 2016; Bouhali et al., 2024; Cachia et al., 2018).

Consistent with this literature, we observed similar prevalence and spatial location of posterior and anterior OTS interruptions, irrespective of study site, sample size, and diagnosis (dyslexia vs control). Within our study and across the literature, the prevalence of the lhpOTS interruption is comparable across children and adults with typical reading skills, children with dyslexia, ex-literate adults, and illiterate adults (Cachia et al., 2018). The lhaOTS interruption appears even more frequently, appearing in ∼70% of individuals in our samples and in the fsaverage template brain (FreeSurfer, Dale et al., 1999). Thus, OTS interruptions are not anomalous anatomical features but rather common manifestations of cortical folding across the population.

Additionally, OTS morphology varies substantially across individuals. While variability is evident in the binary classification of interruptions, it is more pronounced in the degree of sulcal segmentation captured by continuous measures. This suggests that sulcal interruptions represent graded anatomical features that capture meaningful anatomical variation that binary measures obscure. Even continuous measures, however, likely underestimate the full variability of annectant gyri. The OTS relationship to reading only captures *superficial* expressions of annectant gyri and excludes buried expressions located within the sulcal fundus (Mangin et al., 2019). Future work combining surface and volumetric methods to quantify buried expressions (Cykowski et al., 2008; Hopkins et al., 2010, 2014; McKay et al., 2013; Muellen & Schweizer, 2026) are necessary to capture the full spectrum of anatomical variation relevant for behavior. Nonetheless, superficial interruptions vary in size and extent along a continuum, indicating that OTS folding is flexibly expressed across individuals. This combination of anatomical stability and graded variability provides a strong foundation for detecting reliable population-level relationships between sulcal morphology and reading.

### OTS anatomy is not a biomarker of dyslexia

Although OTS morphology relates to reading ability, this relationship varies across the continuum of reading skills. In typical readers, the association between OTS interruptions is robust and replicable. This may reflect the idea that subtle variation in OTS folding may bias the efficiency or organization of visual word processing, producing graded differences in reading performance. In contrast, this relationship does not extend to children with dyslexia. Thus, OTS morphology is not a universal predictor of reading ability but instead relates to reading variability within specific developmental populations. Consequently, the lhpOTS interruption may index exceptionally good reading skills rather than vulnerability to reading impairment.

This distinction has important implications for interpreting OTS morphology as a potential biomarker. Biomarkers are typically expected to reliably differentiate diagnostic groups, reflect underlying biological risk, or predict future outcomes. OTS morphology satisfies none of these criteria. Sulcal patterns do not reliably distinguish between dyslexic versus control participants. Furthermore, when the educational environment is controlled, the presence of a lhpOTS interruption did not predict responsiveness to reading intervention. Instead, all children with dyslexia who received high-quality, evidence-based intervention exhibited significant improvements in reading, regardless of OTS patterns. While anatomy may exert subtle or delayed influences that emerge over longer timescales (Bouhali et al., 2024), other experience-dependent factors – individualized instruction, school curriculum, home environment, etc.(Christodoulou et al., 2017; Donnelly et al., 2019; Hamilton et al., 2016; Huber et al., 2018; Krafnick et al., 2011) – likely play a larger role in shaping outcomes for dyslexic children.

### The function of the lhpOTS and lhaOTS interruptions and annectant gyri

Like other annectant gyri, the lhpOTS and lhaOTS interruptions are prevalent and stable features of human cortical anatomy, yet their origins and significance remain poorly understood. A classical theory dating back to Cunningham (1890) proposes that cortical folding reflects regional differences in function; cortex with greater functional activity may expand more or faster, producing annectant gyri that coincide with the location of functional regions. Examples include the hand knob in the central sulcus in humans, where hand sensorimotor function coincides with a prominent annectant gyrus (Cykowski et al., 2008; Hopkins et al., 2010, 2014; McKay et al., 2013; Muellen & Schweizer, 2026), and primary cortical folds in newborns linked to later functional specialization (Dubois et al., 2008). In monkeys, annectant gyri of the STS predict the location of multiple face patches (Arcaro et al., 2020), suggesting a close link between functional specialization and folding patterns. Applied to the OTS, this framework suggests that sulcal interruptions may anchor the development of functional regions in VOTC. While the OTS itself emerges early in development, functional specialization for visual words develops later and is experience-dependent. Under this view, OTS interruptions may reflect early anatomical biases that interact with experience to shape the organization of word-selective regions.

A complementary account links annectant gyri to white matter organization. In humans and other primates, short-range u-shaped fiber tracts preferentially terminate in gyral crowns and can connect neighboring regions (Nie et al., 2012; Oishi et al., 2011; Zhang et al., 2014). Empirical work has reported higher densities of local u-shaped fibers in superficial annectant gyri in the STS (Le Guen et al., 2018), central sulcus (Mangin et al., 2019; Skandalakis et al., 2025), and OTS (Bouhali et al., 2024). and suggest that annectant gyri might influence local wiring patterns and bias how information is integrated within VOTC. Thus, annectant gyri and their associated u-fibers might influence local wiring patterns, biasing spatial layout and how information is integrated from functional regions within VOTC (Dubois et al., 2008).

Together, these frameworks raise the possibility that the annectant gyri in the OTS and their respective white matter connections may be involved in the emergence of functional regions specialized for word recognition. Evidence from nearby face- (Arcaro et al., 2020; Weiner et al., 2014) and place-selective regions (Natu et al., 2021) anchored to sulcal landmarks and the VWFA predicted by white matter connections (Grotheer et al., 2021; Saygin et al., 2012) supports this possibility. Future work exploring links between the interruptions, functional regions, and white matter connections will be essential for understanding if the OTS interruptions scaffold the functional organization of the reading network.

### Conclusions

Our findings distinguish neuroanatomical correlates of skilled reading from biomarkers of reading impairment. Although variability in OTS anatomy is linked to reading skills in typical readers, it cannot diagnose dyslexia, explain its severity, or predict capacity to benefit from instruction. Dyslexic children therefore do not simply represent one extreme of a continuous anatomical distribution. Rather, anatomical features associated with high reading proficiency are distinct from the mechanisms that govern difficulties in reading. The underlying anatomy does not limit children’s ability to benefit from targeted interventions or determine learning outcomes. More broadly, these findings highlight the importance of the educational experience and targeted interventions, and motivate an integrative approach examining how cortical folding, white matter, functional specialization, and experience jointly shape the developing reading circuitry.

## Acknowledgments

This work was supported by the National Science Foundation Graduate Research Fellowship Program Grant No. DGE-2146755 awarded to J.K.Y. and NICHD R01-HD095861, R01HD116845 to J.D.Y. funding for data collection and curation was provided by NICHD R01-HD095861, R01HD116845 (J.D.Y.) and NIH RO1EY022318, RO1EY023915 (Grill-Spector). We thank Dr. Maya Yablonski, Mia Jimenez, and Hannah Stone, and former members of the Brain Development and Education Lab and Stanford Vision and Perception Neuroscience Lab for assistance with data collection, and the families who participated in the study. We also thank Dr. Vaidehi Natu and Dr. Kalanit Grill-Spector for input on the manuscript.

