## Supplementary Figures for "Sulcal anatomy of ventral temporal cortex and reading development"

**SUPPLEMENTARY MATERIALS**

**Figure S1. Definitions of OTS components in all subjects.** OTS sulcus (black), anterior OTS gap (coral), posterior OTS gap (turquoise) outlined in the typical and dyslexic participants from the Stanford-I, Seattle-I, Stanford-D, and Seattle-D studies.

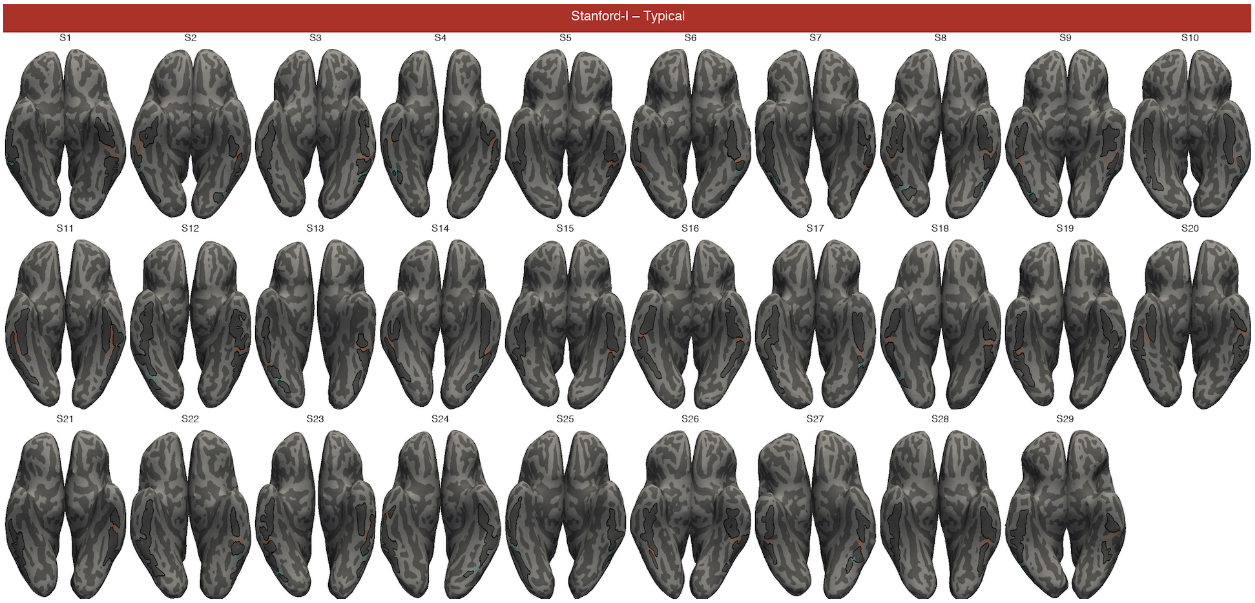

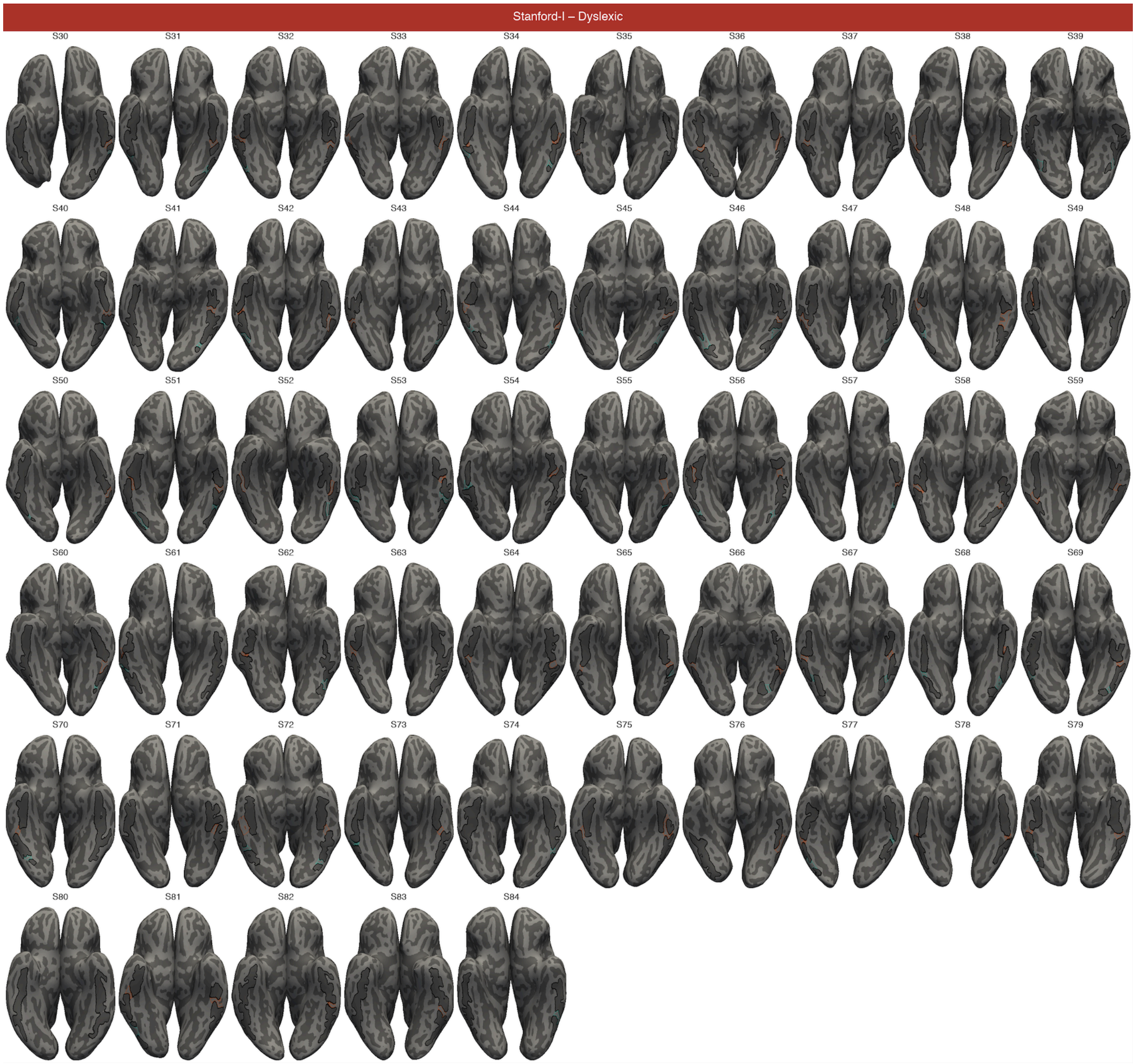
Stanford-I – Dyslexic
S30
S31
S32
S33
S34
S35
S36
S37
S38
S39
S40
S41
S42
S43
S44
S45
S46
S47
S48
S49
S50
S51
S52
S53
S54
S55
S56
S57
S58
S59
S60
S61
S62
S63
S64
S65
S66
S67
S68
S69
S70
S71
S72
S73
S74
S75
S76
S77
S78
S79
S80
S81
S82
S83
S84

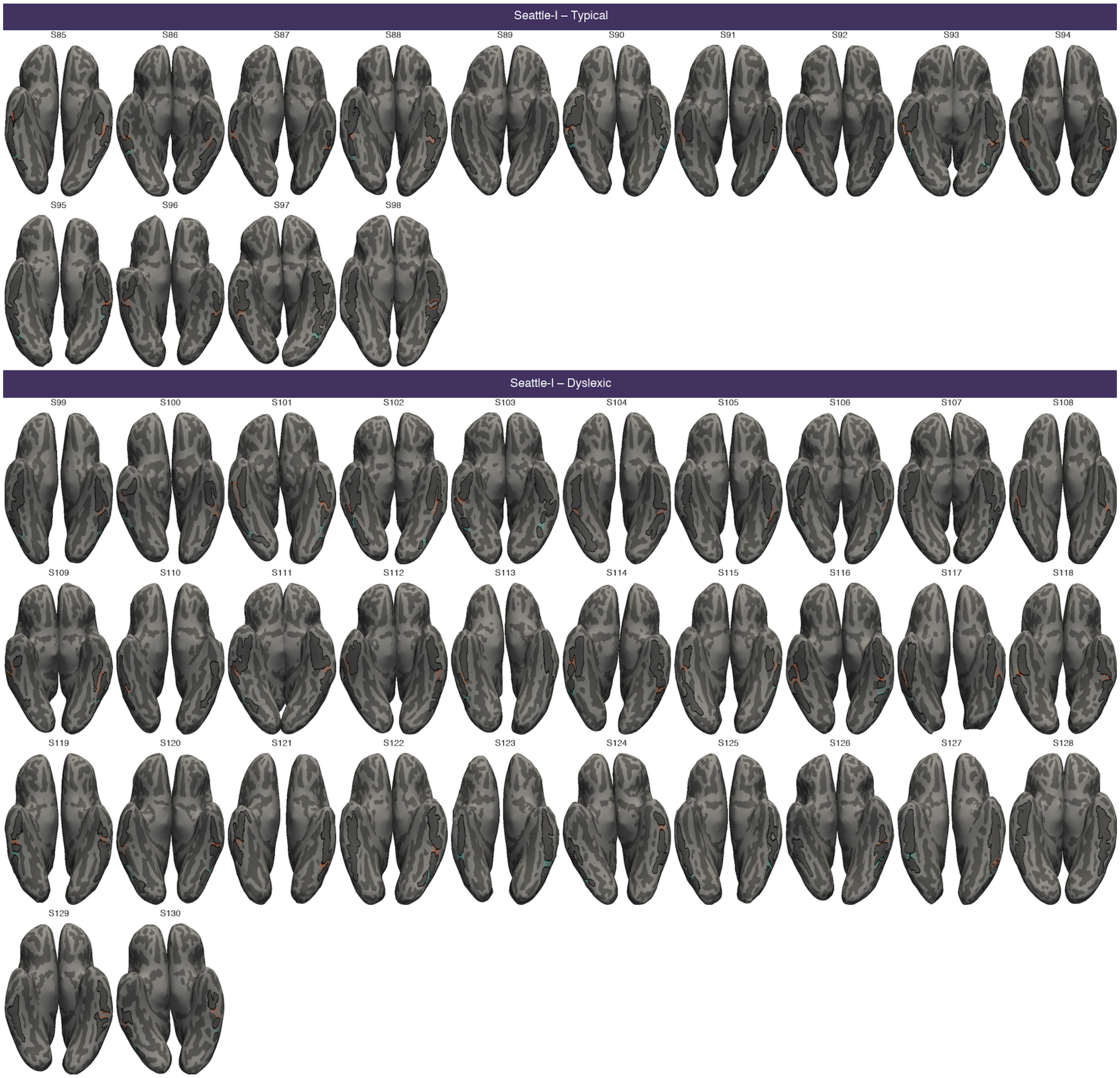
Seattle-I – Typical
S85
S86
S87
S88
S89
S90
S91
S92
S93
S94
S95
S96
S97
S98
Seattle-I – Dyslexic
S99
S100
S101
S102
S103
S104
S105
S106
S107
S108
S109
S110
S111
S112
S113
S114
S115
S116
S117
S118
S119
S120
S121
S122
S123
S124
S125
S126
S127
S128
S129
S130

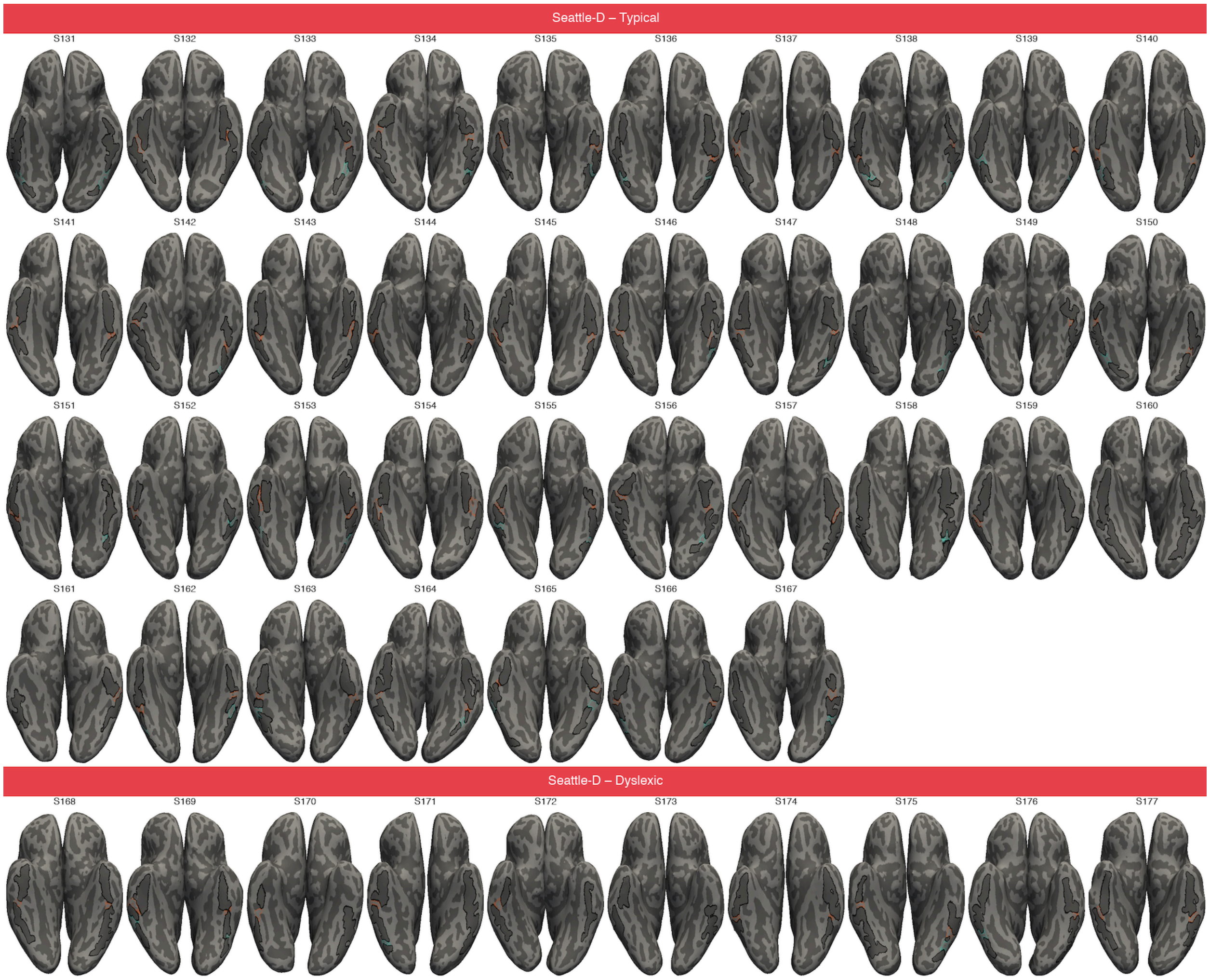
Seattle-D – Typical
S131
S132
S133
S134
S135
S136
S137
S138
S139
S140
S141
S142
S143
S144
S145
S146
S147
S148
S149
S150
S151
S152
S153
S154
S155
S156
S157
S158
S159
S160
S161
S162
S163
S164
S165
S166
S167
Seattle-D – Dyslexic
S168
S169
S170
S171
S172
S173
S174
S175
S176
S177

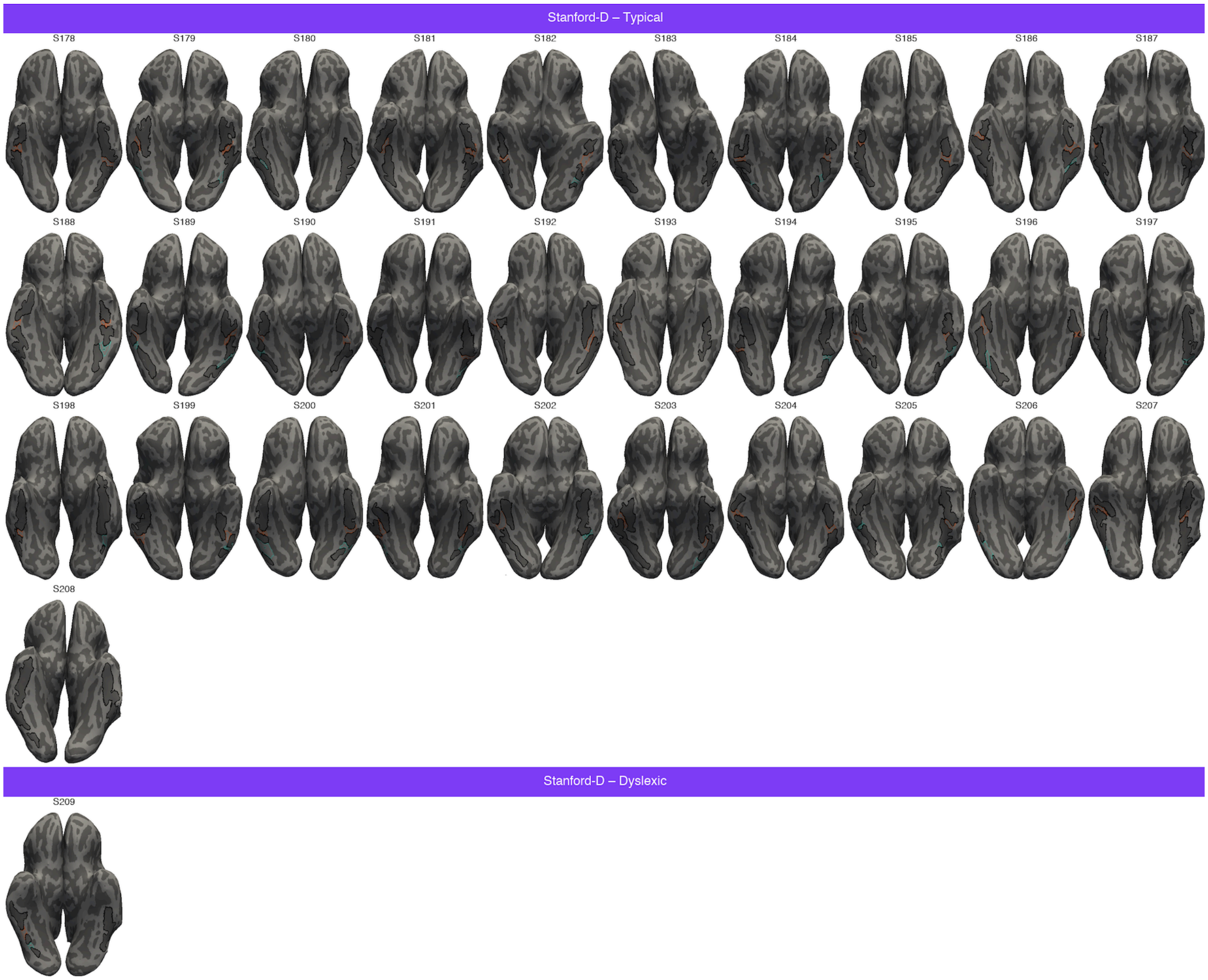

Stanford-D – Typical
S178
S179
S180
S181
S182
S183
S184
S185
S186
S187
S188
S189
S190
S191
S192
S193
S194
S195
S196
S197
S198
S199
S200
S201
S202
S203
S204
S205
S206
S207
S208
Stanford-D – Dyslexic
S209

**Figure S2. Definitions of VOTC sulci in all subjects.** Occipital temporal sulcus (OTS; black), mid fusiform sulcus (mfs; red), collateral sulcus (CoS; green), anterior transverse CoS (atCoS; white), posterior transverse CoS (ptCoS; blue) outlined in the typical and dyslexic participants from the Stanford-I, Seattle-I, Stanford-D, and Seattle-D studies.

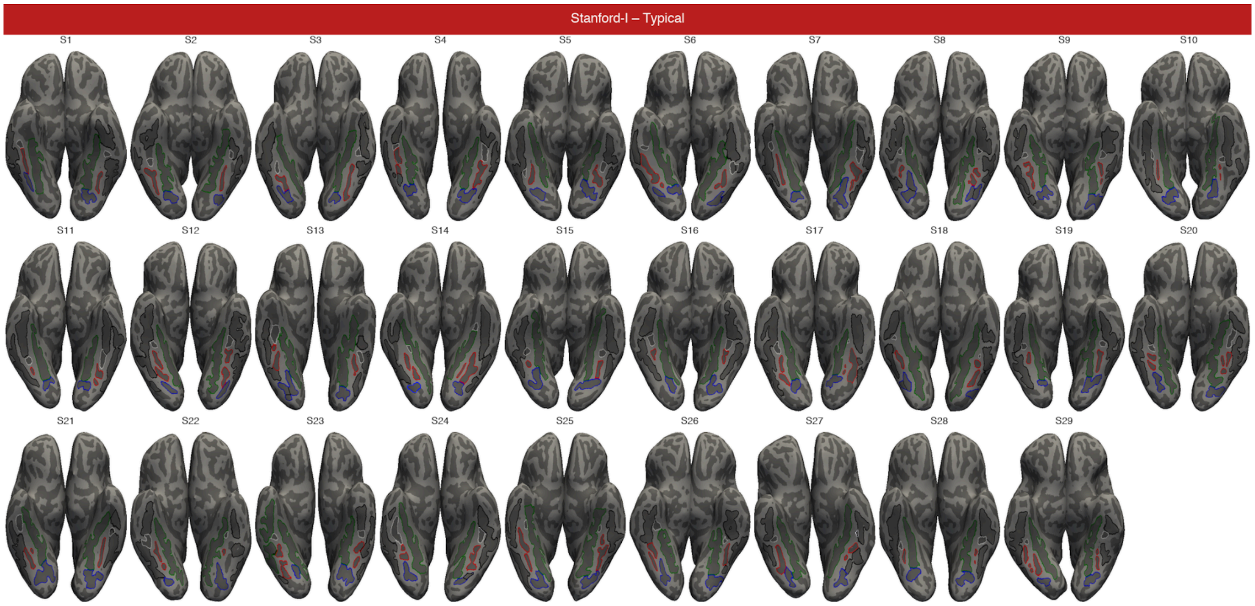

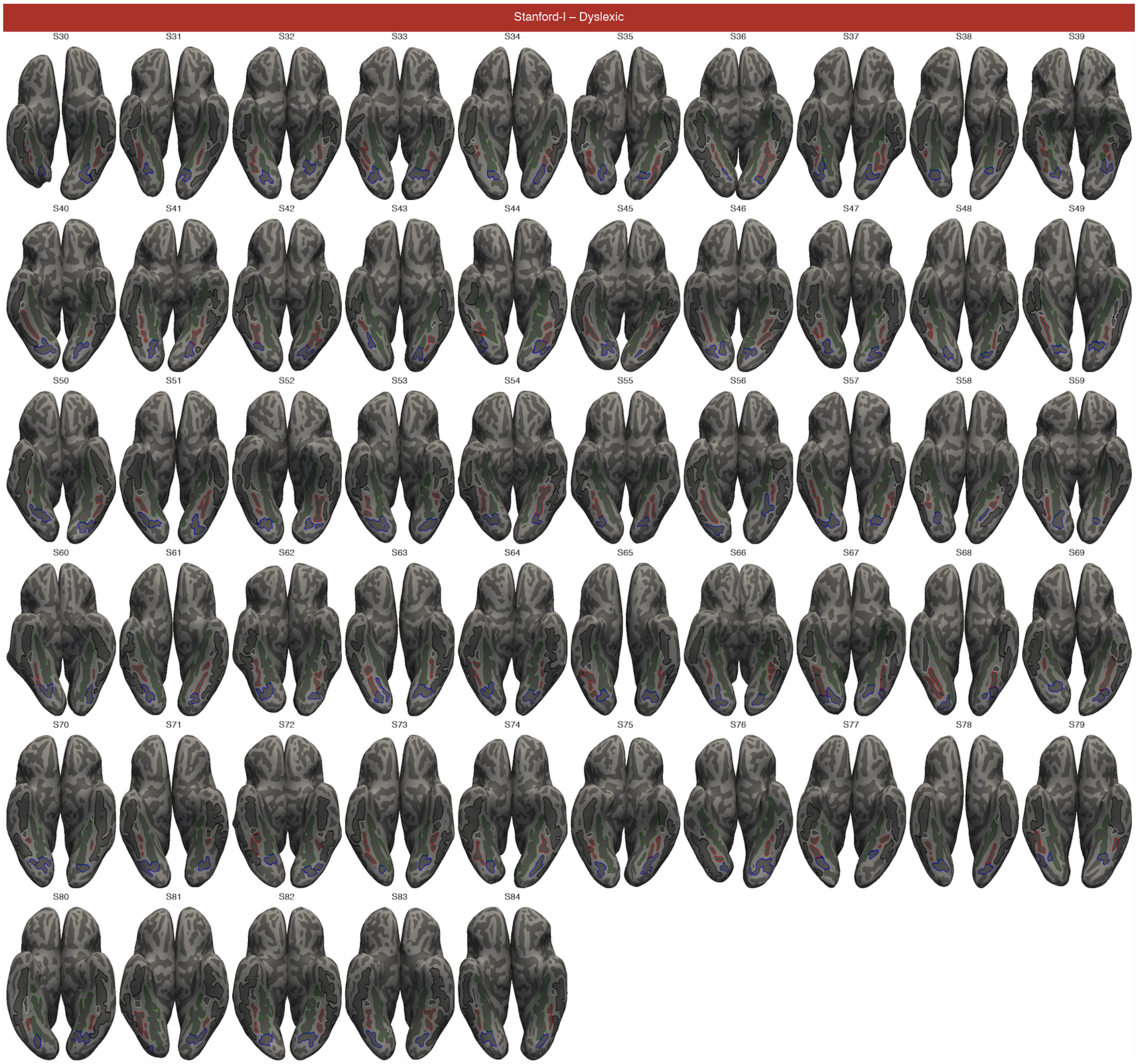
Stanford-I – Dyslexic
S30
S31
S32
S33
S34
S35
S36
S37
S38
S39
S40
S41
S42
S43
S44
S45
S46
S47
S48
S49
S50
S51
S52
S53
S54
S55
S56
S57
S58
S59
S60
S61
S62
S63
S64
S65
S66
S67
S68
S69
S70
S71
S72
S73
S74
S75
S76
S77
S78
S79
S80
S81
S82
S83
S84

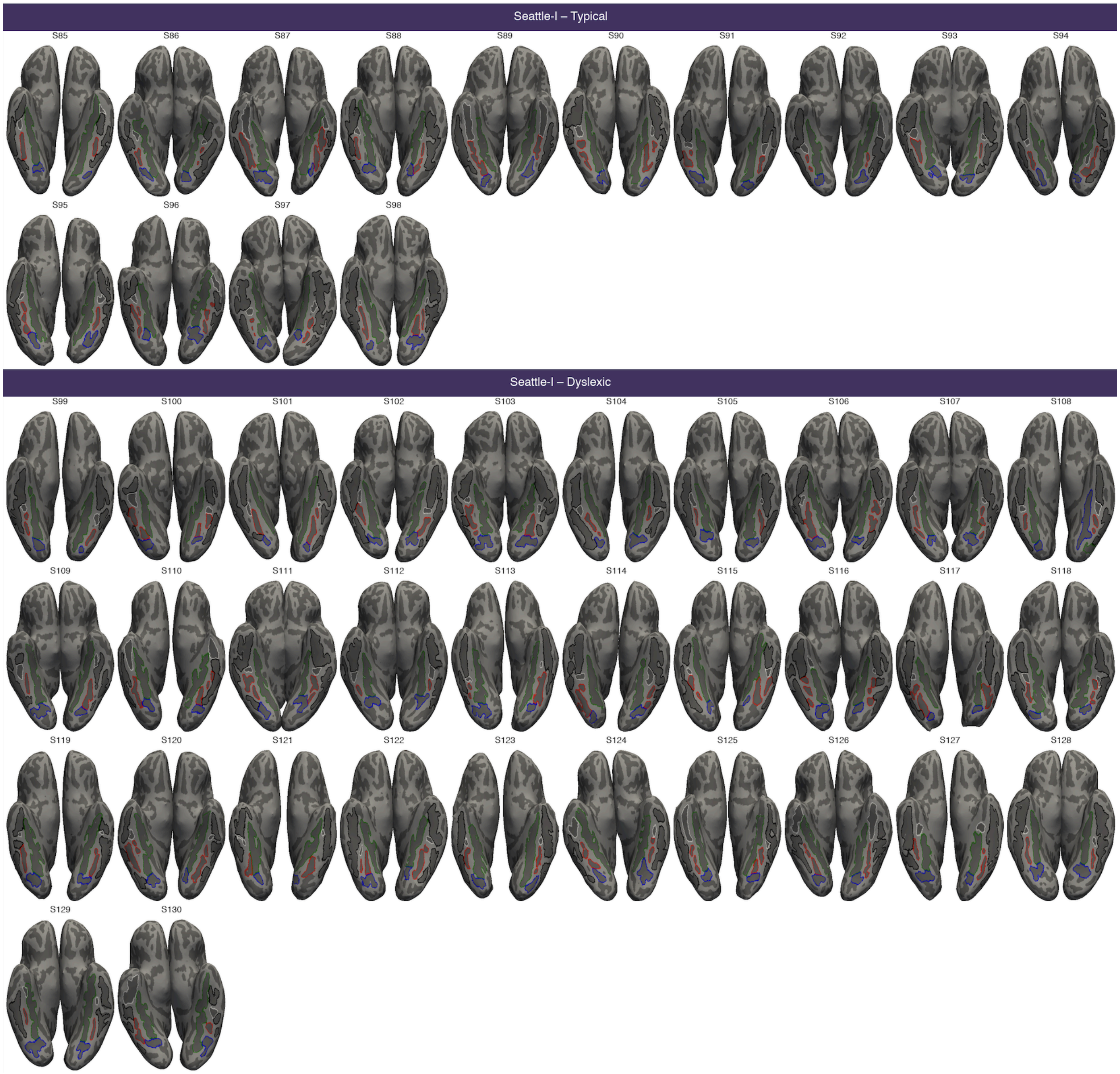

Seattle-I – Typical
S85
S86
S87
S88
S89
S90
S91
S92
S93
S94
S95
S96
S97
S98
Seattle-I – Dyslexic
S99
S100
S101
S102
S103
S104
S105
S106
S107
S108
S109
S110
S111
S112
S113
S114
S115
S116
S117
S118
S119
S120
S121
S122
S123
S124
S125
S126
S127
S128
S129
S130

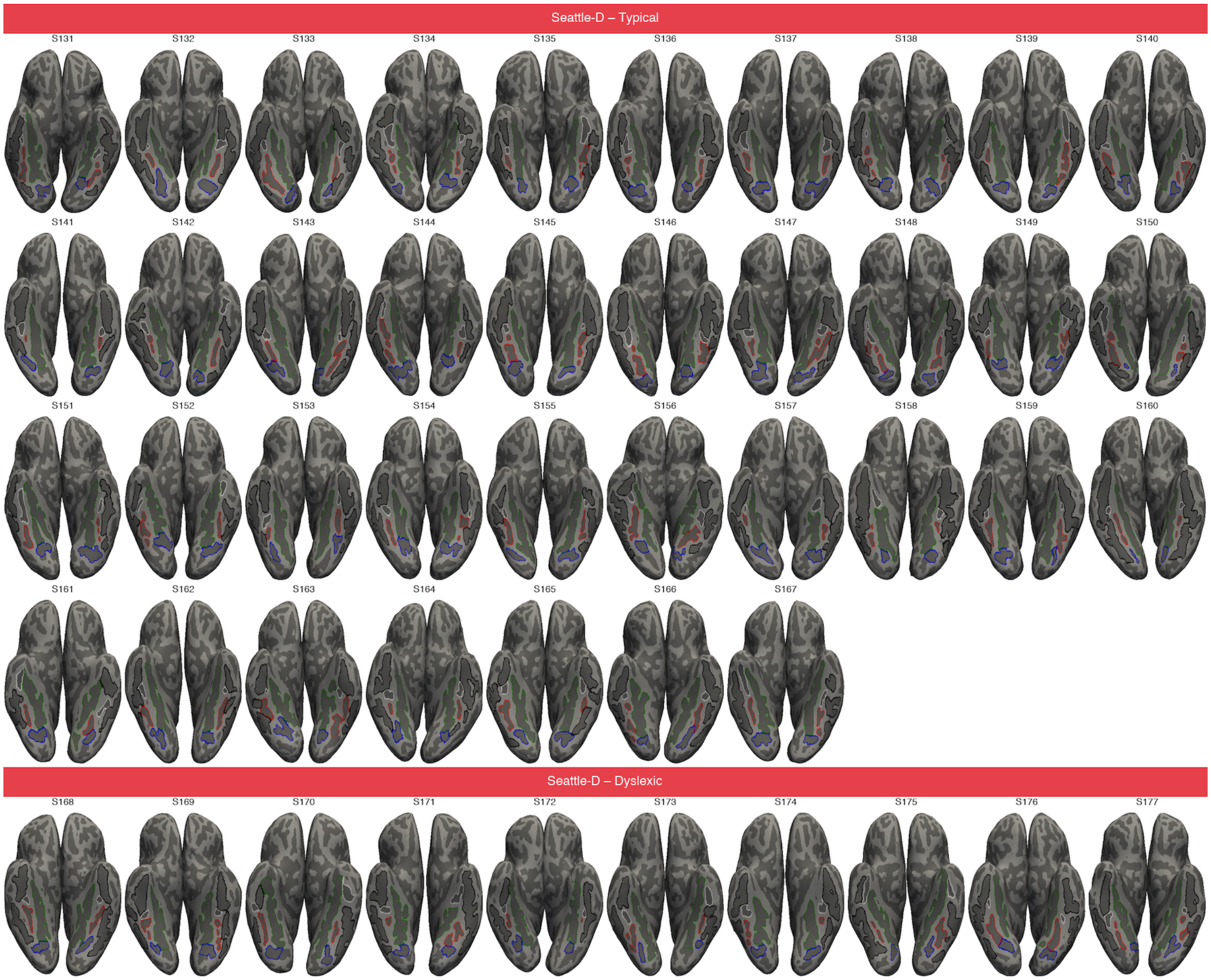
Seattle-D – Typical
S131
S132
S133
S134
S135
S136
S137
S138
S139
S140
S141
S142
S143
S144
S145
S146
S147
S148
S149
S150
S151
S152
S153
S154
S155
S156
S157
S158
S159
S160
S161
S162
S163
S164
S165
S166
S167
Seattle-D – Dyslexic
S168
S169
S170
S171
S172
S173
S174
S175
S176
S177

Stanford-D – Typical

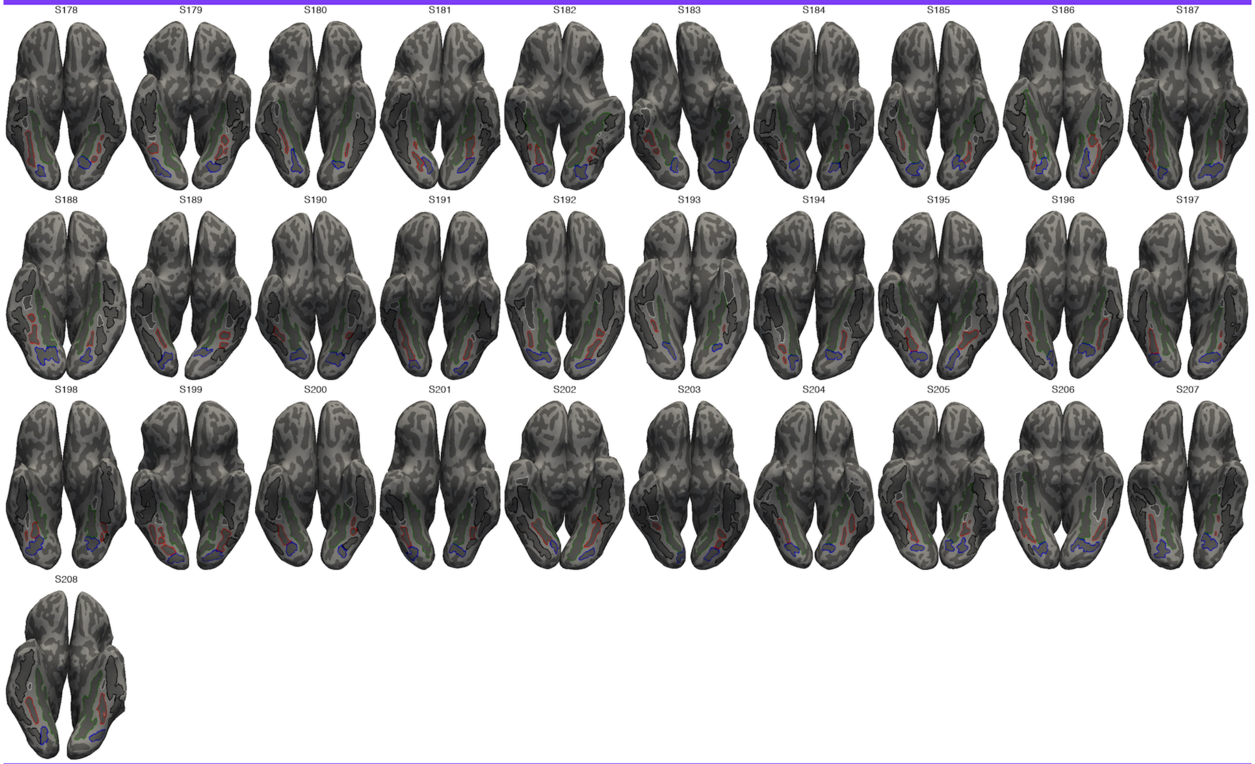


Stanford-D – Dyslexic

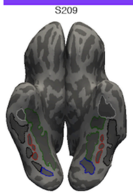
